# Exploring Effective Ingredients and Potential Mechanisms of Stem Cell-derived *Dendrobium Officinale* Extract in Treatment of Allergic Rhinitis

**DOI:** 10.1101/2025.04.18.649518

**Authors:** Jiangning Li, Jie Zhang, Ken Cheng

**Affiliations:** Xi’an Jiaotong-Liverpool University, Wisdom Lake Academy of Pharmacy, Suzhou, Jiangsu, China

**Author notes:** Correspondence: Jie Zhang, Tel (optional), Fax (optional), Email (mandatory). Correspondence: Ken Cheng, Tel (optional), Fax (optional), Email (mandatory).

**Keywords:** *Dendrobium Officinale*, allergic rhinitis, metabolomic, network pharmacology

## Abstract

**Background:** Dendrobium Officinale is a rare Chinese herbal medicine, and its stem has various medical effects. Currently, the sprouts of Dendrobium Officinale can be mass-produced by stem-cell technology, but its biological active ingredients and medical effects are unknown. The therapeutic ability of the stem cell-originated *Dendrobium Officinale* (SDO) for allergic rhinitis (AR) was demonstrated in the preliminary experiment on mice.

**Aim:** Metabolomics methods were employed to analyze the differences in metabolomics between SDO and commercial *Dendrobium officinale* stem (COM) water extracts, and verify the bioactive ingredients related to treating AR. A network pharmacology-based strategy was applied to investigate potential therapeutic targets and mechanisms of SDO against AR. Molecular docking was performed for validation.

**Materials and Methods:** This study used HPLC-MS/MS-based metabolomics to analyze the metabolomic differences between SDO and COM water extract, and identified some active ingredients related to AR treatment. Network pharmacology was used to screen the potential targets for SDO treats AR, establish protein-protein interaction (PPI) networks, and perform enrichment analysis (KEGG/GO) to explore the signalling pathways. Molecular docking was applied to further verify the result.

**Results:** The metabolomic differences between SDO and COM were demonstrated.2053 differential metabolites were found and 6 bioactive ingredients related to AR treatment were verified. 121 common targets of SDO and AR were identified in network pharmacology, the top ten core targets were screened based on the PPI network, and 7 possible therapeutic mechanisms that SDO treat AR were speculated. Molecular docking proved that each core target combined well with the validated bioactive ingredients.

**Conclusion:** This study compared the metabolomes of SDO and COM, and explored the bioactive components, therapeutic targets, and potential mechanisms of SDO in treating AR. It explored the medical potential of SDO for the first time and provided a promising direction for the large-scale medical application of *Dendrobium Officinale*.

## Introduction

Dendrobium Officinale is a harmless and non-toxic herbal medicine, which is often used to treat gastrointestinal diseases.^1^ It has been used as the nutraceutical food or drink material and is recognized as the best tonic in the dendrobium species.^2^ Current researches show that the Dendrobium Officinale and its extracts have multiple medical effects such as hypoglycemic, antihypertensive, hypolipidemic, antioxidant, anti-tumour, anti-fatigue, enhancing immunity, enhancing vision and gastrointestinal health.^2,3^ According to Wang et al., studying the metabolomes of Dendrobium officinale is important for exploring its medicinal properties.^4^ Currently, some researchers have focused on the metabolomic of Dendrobium officinale. An experimental group in Kunming (China) identified 133 metabolites containing nitrogen in Dendrobium officinale.^5^101 volatile components were identified in Dendrobium officinale from 4 regions.^6^ 411 secondary metabolites of Dendrobium officinale have been identified by Luo et al.^7^ However, in different plant parts or producing regions or, the metabolomes of Dendrobium officinale have differences.^6,8^ The metabolomic differences may lead to changes in treatment effects.

Due to the special growth pattern requiring root symbiotic bacteria for seed germination, the natural reproduction rate of wild Dendrobium Officinale is low.^9^ The Dendrobium Officinale stems are commonly utilized for medication. However, due to the scarce production and high prices, it is difficult to apply on a large scale.^9^ Research has found that there is a technique that helps achieve large-scale production of Dendrobium Officinale buds. Zhang proposed a technique that expands Dendrobium Officinale stem cells through in vitro large-scale suspension culture to achieve the mass production of the Dendrobium Officinale sprouts.^10^ It demonstrates that, the stem cell-originated Dendrobium Officinale (SDO) may already have the foundation to be utilized in bulk quantities. However, most of the studies about Dendrobium Officinale focus on the stems or leaves, but metabolites and medicinal effects of the Dendrobium Officinale sprouts are still unknown.^2^

Rhinitis, including allergic rhinitis (AR), is the most common upper respiratory tract inflammatory disease.^11^ The incidence rate of AR has raised rapidly in the past few decades and is often accompanied by complications such as allergic asthma.^12^ Currently, conventional drugs for treating allergic rhinitis mainly aim to alleviate symptoms, and their side effects are significant such as drowsiness and arrhythmia.^13^ Therefore, new drugs targeting the mechanisms of allergic rhinitis with fewer side effects need to be developed. According to Ma et al., the Dendrobium Officinale could reduce the inflammatory cells’ infiltration and the inflammatory factors’ levels, repair and protect the gastric mucosa.^14^ Rhinitis and gastric mucosal damage are both inflammations in the mucosal area, and it may be inferred that Dendrobium Officinale probably has some therapeutic effects on allergic rhinitis.

To investigate whether Dendrobium Officinale has a therapeutic effect on AR, we designed a preliminary experiment using SDO water extract to treat OVA-induced allergic rhinitis in mice was conducted. The processes of the preliminary experiment are shown in Figure 1, and the therapeutic effect was evaluated through behavioral experiments.

**Figure 1.**
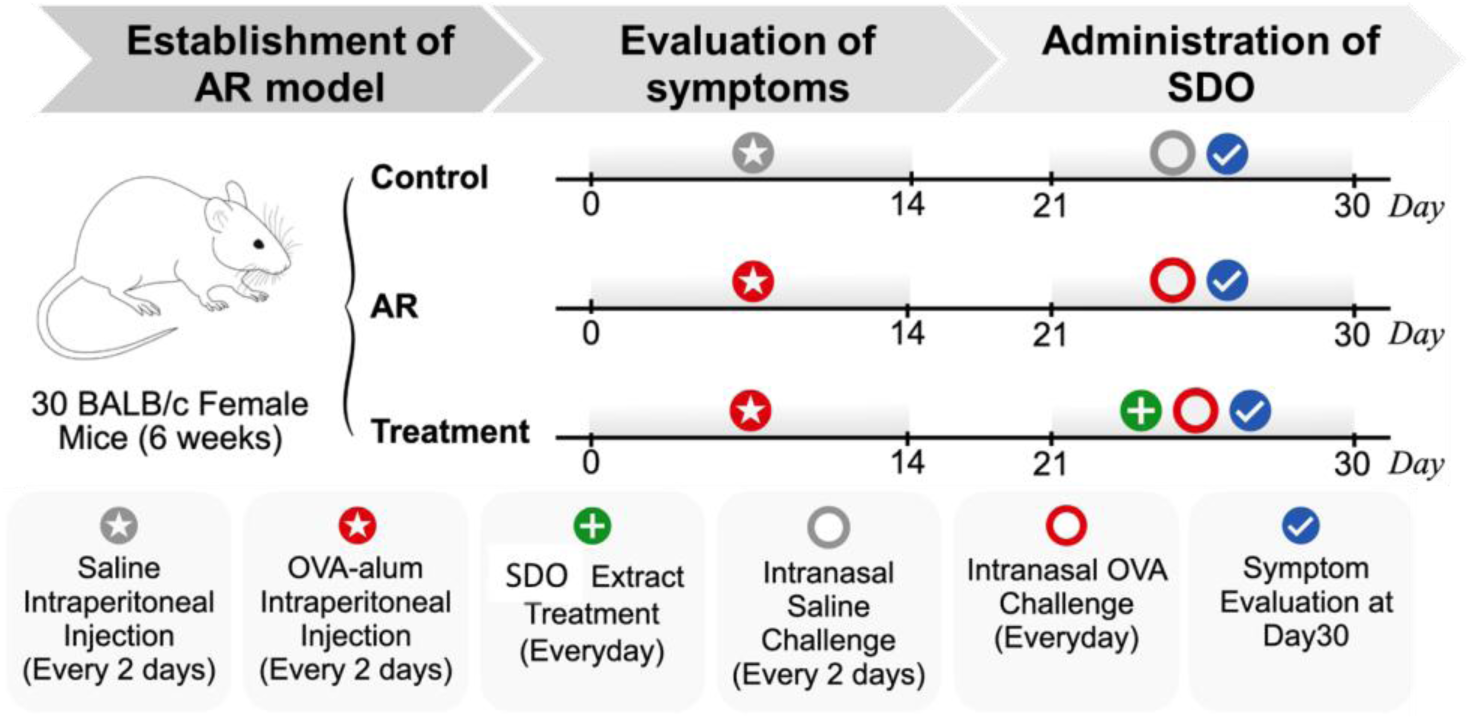
The process of the pilot study on the treatment of OVA-induced allergic rhinitis in mice by the stem cell-originated Dendrobium Officinale (SDO) water extract.

According to figure 2, scratching nose times and sneezing times in the treatment group mice was significantly less frequently than the AR group mice, the symptoms of AR were significantly relieved after receiving the SDO water extract. But the treatment group still has some symptoms compared to the control group, it means the mice could not recover completely at this dose. In summary, the result of the preliminary experiment proves that SDO water extract has a certain effect on the AR treatment. Still, the treatment mechanism has been unclear.

**Figure 2.**
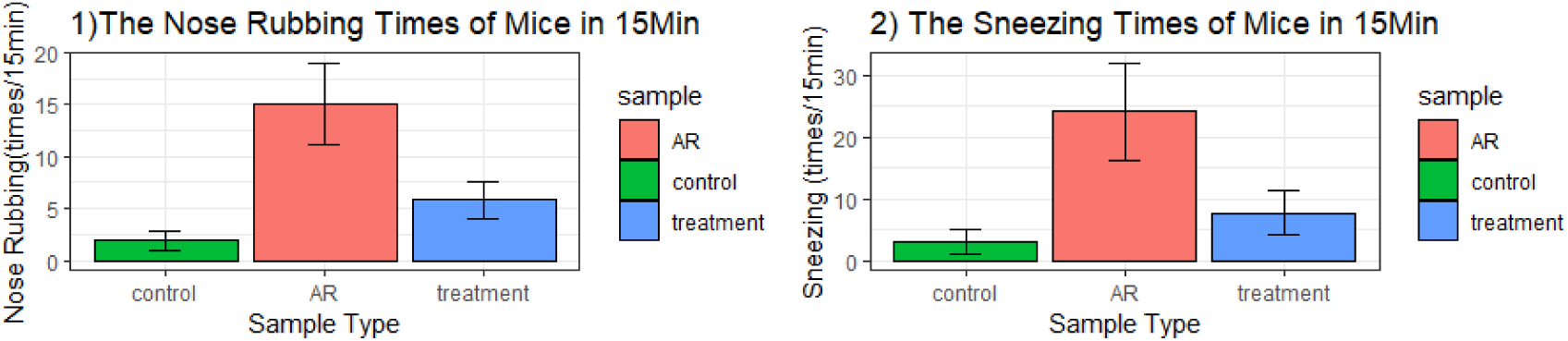
The sneezing (1) & nose rubbing (2) frequencies of mice within 15 minutes after the intranasal OVA in day21-30, expressed as mean ± SD. The post hoc test is Game-Howell test, and p.adj < 0.05

Plant metabolites are the source of many natural products with pharmacological activities, and metabolomics has been utilized in analyzing herbal components extensively.^15^ The action mechanism of traditional Chinese medicine is similar to network pharmacology. Both of them are holistic, comprehensive and systemic.^16^ Therefore, network pharmacology is appropriate for exploring Chinese herbal medicine’s pharmacological mechanism and analyzing and constructing the network models of drug effects.^16^ To explore the application potential of SDO, especially in the treatment of allergic rhinitis, we used metabolomics to analyze the metabolite differences between SDO and commercial Dendrobium Officinale (COM) water extracts, validated some metabolites in SDO related to the therapy of allergic rhinitis, and investigated the possible mechanisms of SDO water extract treatment for allergic rhinitis through network pharmacology.

## Material and methods

### Herbal Materials, Reagents and Chemicals

The stem cell-originated Dendrobium Officinale (SDO) extract was obtained from Suzhou Planterbio Stem-cell Technology Co., Ltd., the commercial Dendrobium Officinale (COM) extract was purchased from Suzhou Tiara Biotech Co. Ltd.

Methanol (LC/MS grade) was purchased from Macklin Inc. Formic acid and acetonitrile (LC/MS grade) were purchased from Thermo Fisher Scientific Inc. Water (LC/MS grade) was purchased from Alfa Aesar.

(S)- Equal, 4-phenylbutyric acid, fumalic acid, 4-coumaric acid, urocanic acid and azelaic acid were purchased from the Glpbio Technology. Allantoin, nodakenin and 4-methyl-5-thiazoleethanol were purchased from Shanghai Aladdin Biochemical Technology Co., Ltd. The purities of all the above chemicals are more than 98%.

### Metabolomics Analysis

#### Sample and Standard Preparation

The water extracts of stem cell-originated dendrobium officinale sprout extract and traditional dendrobium officinale extract were extracted and dissolved into 1mg/ml by PBS. Then the two water extracts of 1mg/ml were dissolved into 30 ppm by 50% acetonitrile.

4-phenylbutyric acid, fumalic acid, 4-coumaric acid, urocanic acid, nodakenin and 4-methyl-5-thiazoleethanol were dissolved by 50% methanol into 1mg/ml, and then dissolved into 5 and 1 ppm by 50% acetonitrile. (S)- Equal, azelaic acid and allantoin were dissolved into 5 and 1 ppm by 50% acetonitrile.

#### Chromatographic Conditions

The chromatographic separation used Thermo Hypersil GOLD aQ columin (2.1 mm ×100 mm, ThermoFisher) as the chromatographic column. The mobile phase A is ACN (acetonitrile) with 0.1% formic acid, and mobile phase B is H2O with 0.1% formic acid. The gradient of the mobile phase was as followed: 0-2 min, 5% A; 2-42 min, 5%-95% A; 42-47min, 95% A; 47.1-50 min,5% A. The Flow rate was 0.3 mL/min, the injection volume was 5 μL, and column temperature was 40°C.

#### Mass Spectrum Conditions

HPLC-MS/MS was UPLC-Q Executive plus and performed by the Ultimate 3000 HPLC-QE plus MSMS Thermo. The ESI ionization was both positive and negative. The scanning range (m/z) was 100 - 1500; scan mode was Full MS / ddMS2; the spray voltage was 3.2kV (-)/3.5kV (+); the ion source temperature was 320 °C; the Aux gas (N2) pressure is 10 arb; the sheath gas (N2) pressure is 35 arb; the MS max IT was 100 ms; the AGC target is 1×106. The raw data were processed in Compound Discovery software in which the mzcloud and OTCML databases were applied for metabolite identification. The credibility of the identified compound names of variables was determined based on the matching results of Predicted Compositions, mzCloud, and ChemSpider databases. When the matching result of the mzCloud database is "full match" or the matching results of Predicted Compositions and ChemSpider databases are both "full match", the compound name is considered credible.

#### HPLC-QToF-MSMS Analysis for Validation

The analysis was carried out by HPLC 1290-QToF MSMS 6530C (Agilent). Chromatographic conditions was 2.2.1 The gas temp was 350℃ and the dry gas was10 l/min. The nebulizer was 45 psi. The sheath gas flow was 12 l/min and the sheath gas temp was 400°C. The Vcap was 4000 V; the capillary was 0.068 μA. The fragmentor was 175V and the skimmer: 65V. The detection window is 20 ppm. The single m/z expansion when extracting the chromatograms is +/- 20 ppm. The experiment was repeated at three times.

#### MS Data Processing

The obtained metabolic data (.raw files) was converted into CSV files by Compound Discovery software. The support vector regression (SVR) normalization was used for the data standardization, and the variables with coefficients of variation (CV) < 30% in the QC groups were reserved for data analysis. The function used for SVR normalization is from the R package “MetNormalizer”.^17^ P-value, FDR, fold change (mean area of SDO: mean area of COM) and OPLS-DA were used to screen the metabolic differences. Chemicals related to the treatment of allergic rhinitis or inflammation were selected among variables that satisfied p value<0.05 and fold change>2 or VIP>1. All statistical analyses were conducted using R language, R version is 4.1.1.

### Network Pharmacology

#### Target Analysis

The information of L-Arginine, urocanic acid, 4-methyl-5-thiazoleethanol, allantoin, 4-phenylbutyric acid and azelaic acid was obtained from the PubChem database. And the information was utilized to search the drug targets of these substances from PharmMapper (top 100), Swiss Target Prediction database (p>0) and Similarity Ensemble Approach database (MaxTc≥0.4)^17,18,19^. The allergic rhinitis-related targets were obtained from OMIM, Gene Cards, Drug Bank, PharmGKB and Therapeutic target database.^21,22,23,24,25^ Then the intersection of the drug target and the allergic rhinitis target were obtained as the possible targets that Dendrobium Officinale treats allergic rhinitis.

#### The Protein–protein Interaction (PPI) Network Construction

The gene symbols of intersecting targets were uploaded to the STRING database, and the organization was set as "Homo sapiens" with the medium confidence set to high confidence (>0.7).^26^ The short tabular text of the PPI network was downloaded and imported into the Cytoscape 3.9.1 to generate the PPI network of treatment.

#### Screening Core Targets Based on the Topological Structure of PPI Network

Then the short tabular text of the PPI network was uploaded into Cytoscape 3.9.1 for the network topology analysis. The MCC algorithm of the cytoHubba plug-in was utilized to calculate the values of targets.^27^ The targets with the score of top 10 were selected as the core targets for further analysis.

#### Molecular Docking

The molecular structures of validated chemicals were obtained from the PubChem database (https://www.rcsb.org/). The protein crystal structures of the core targets were downloaded from the RCSB PDB database (https://www.rcsb.org/). PyMol was used to separate the ligand of proteins, remove water molecules and add hydrogen atoms. CB dock2 was used to perform blind docking to obtain the affinity of the docking conformation.^28^ The docking configurations were visualized by PyMol.

## Results

### Metabolomic Differences Analysis between SDO and COM

#### Macroscopic Differences in Metabolomics

Compared to PLS-DA and PCA, OPLS-DA is more sensitive and can better filter variables with weaker correlations, maximize inter group differences, and more effectively extract main information.^29^ To understand the metabolomes of SDO and COM at a macro level, OPLS-DA was used to construct classification prediction models on the positive and negative ionization result datasets from HPLC-MS/MS respectively. According to figure 3 (a2 and b2), the values of R2Y and Q2Y of the two models established based on dataset of positive or negative ionization are all not greater than the true value (all scatter points are lower than the horizontal line), so neither model has the overfitting phenomenon. The Q2Y values of the two models are 0.991 (POS) and 0.95 (NEG). Both the Q2Y values close to 1, so the two OPLS-DA models are stable and reliable.

**Figure 3.**
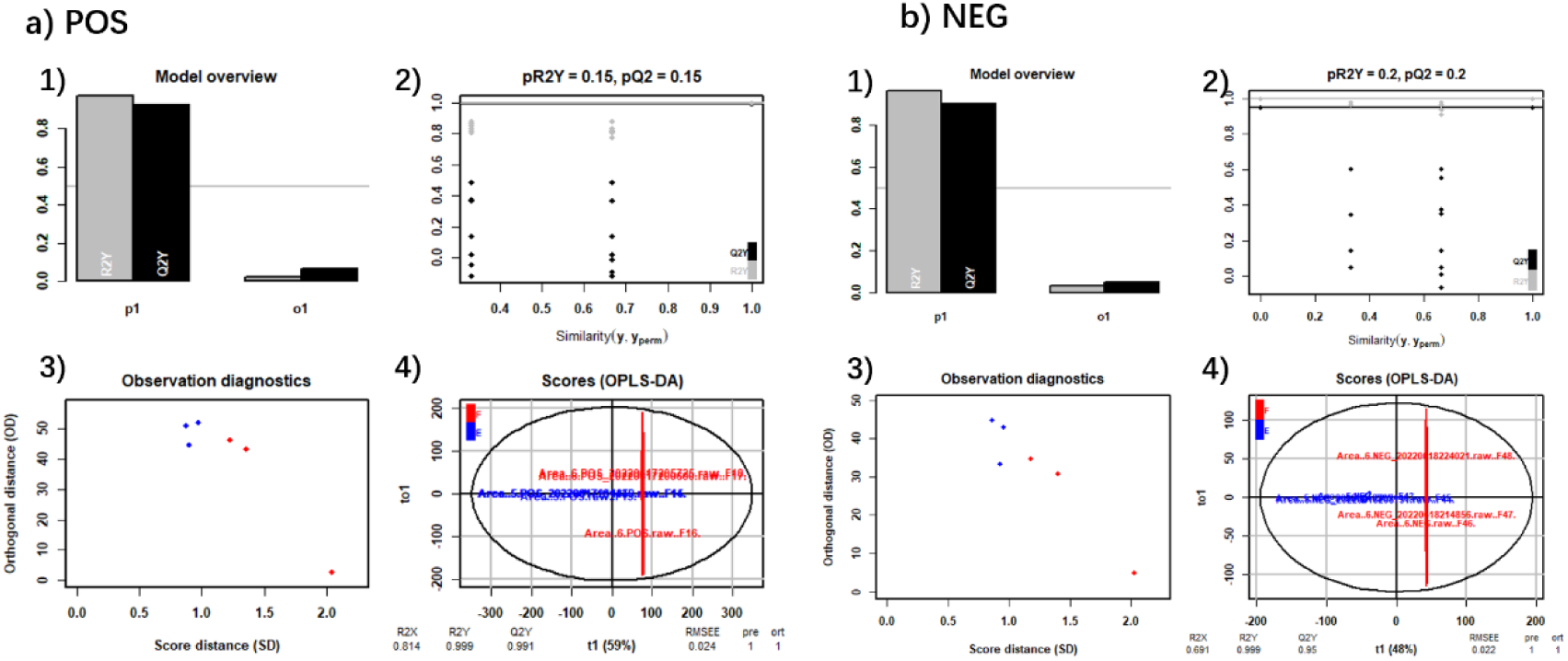
OPLS-DA results. POS: data is from the positive ionization result. NEG: data is from the negative ionization result. In the OPLS-DA score chart, blue represents COM and red represents SDO.

According to the two score plots of the OPLS-DA models (Figure3 a4&b4), the water extract samples of SDO and COM are significantly separated in the two plots, indicating that SDO and COM have observable metabolomic differences according to both the HPLC-MS/MS results from positive and negative ionization forms. The VIP values of each variable calculated by the OPLS-DA model were extracted for further differences analysis of each metabolite, where metabolites with VIP value larger than 1 were considered having the expression difference between the two groups.

#### Specific Metabolite Differences Analysis

Total 23362 metabolites (POS: 16987; NEG: 6375) were obtained by the HPLC-MS/MS. Among them, 20052 variables had credible names (POS: 14455; NEG: 5597), and 16778 metabolites (POS: 12132; NEG: 4646) had CV < 30%. The metabolites whose FDR < 0.05 and the absolute value of log2FC > 1 are considered as having expression differences. According to Figure 4, in the result of positive ionization, most metabolites were downregulated in SDO water extract compared to COM water extract, while in the negative result, there were more metabolites upregulated in the SDO sample compared to the COM sample.

**Figure 4.**
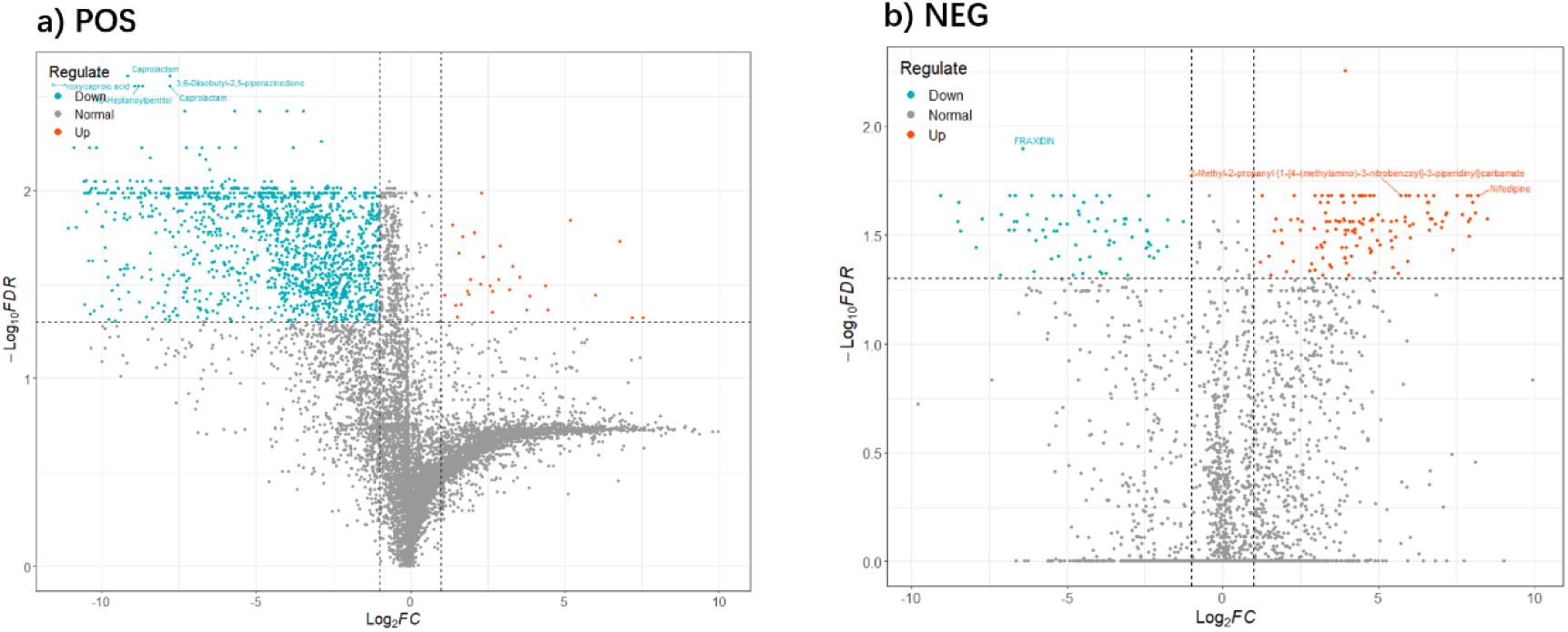
The volcano plots of the metabolites. POS: plotting based on positive ionization results. NEG: according to negative ionization results.

The Venn plot (Figure 5) shows that a total of 175 metabolites were upregulated in SDO samples compared to COM samples, while the number of downregulated samples was 1878, approximately ten times the number of upregulated samples. It indicates that SDO may be more enriched in the expression of a few metabolites.

**Figure 5.**
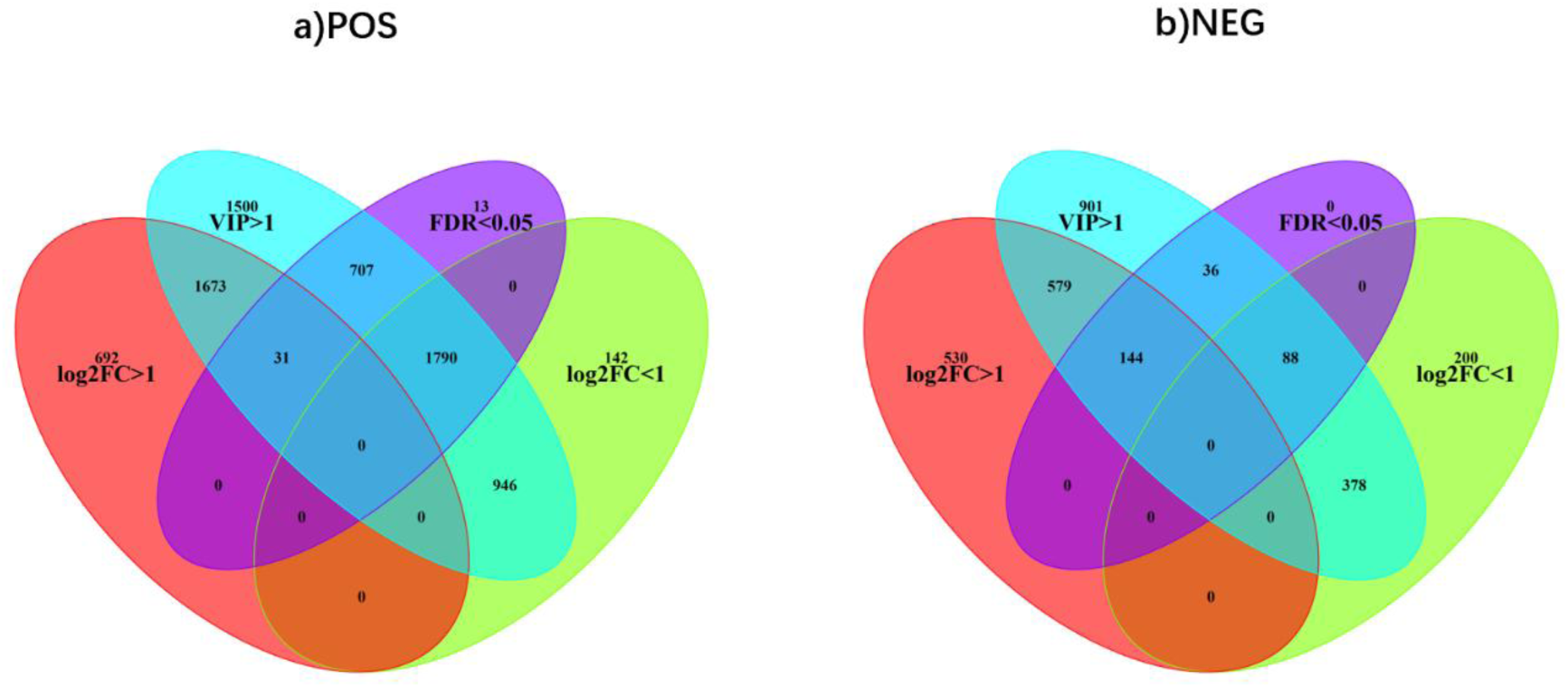
the Venn diagrams of metabolites. POS: the metabolite data is from the positive ionization result. NEG: the metabolite data is from the negative ionization result.

The top five compounds (FDR > 0.05) with the largest difference in fold change (i.e. the largest absolute value of fold change) in the positive or negative ionization HPLC-MS/MS results are shown in Table 1 and 2 respectively.

**Table 1.** The top five metabolites with the highest expression differences in positive ionization results.

| Name | Formula | m/z | rt | P.Value | log<sub>2</sub>FC | FDR |
| --- | --- | --- | --- | --- | --- | --- |
| [4-(Dimethylamino)-2,6-pyridinediyl]dimethanol | C<sub>9</sub>H<sub>14</sub>N<sub>2</sub> O<sub>2</sub> | 458.235 | 11.244 | 3.52E-08 | -11.0596 | 0.015889 |
| L-Ethionine | C<sub>6</sub>H<sub>13</sub>NO<sub>2</sub>S | 502.2616 | 11.732 | 4.81E-07 | -10.8697 | 0.005942 |
| 6-(2-Oxiranylmethyl)-<br>1,2,3,4,5,11-hexahydro-9H-cyclohepta[b]quinolin-9-one | C<sub>17</sub>H<sub>19</sub>NO<sub>2</sub> | 458.2353 | 11.253 | 4.40E-08 | -10.8141 | 0.015838 |
| 6-Hydroxy-2-(2-phenylethyl) chromone | C<sub>17</sub>H<sub>14</sub>O<sub>3</sub> | 477.2312 | 11.406 | 8.47E-06 | -10.5219 | 0.009023 |
| Ethyl (2E)-3-<br>(methylamino)-2-butenoate | C<sub>7</sub>H<sub>13</sub>NO<sub>2</sub> | 521.256 | 11.917 | 3.61E-10 | -10.5168 | 0.010393 |

**Table 2.** The top five metabolites with the highest expression differences in negative ionization form.

| Name | Formula | mz | rt | P.Value | log<sub>2</sub>FC | FDR |
| --- | --- | --- | --- | --- | --- | --- |
| 2-Amino-6-(2,4-dichlorobenzoyl)-1,5,6,7-tetrahydro-4H-pyrrolo[3,4-d]pyrimidin-4-one | C<sub>13</sub>H<sub>10</sub>Cl<sub>2</sub>N<sub>4</sub><br>O<sub>2</sub> | 394.2094 | 9.069 | 6.71E-08 | -9.78207 | 0.189943 |
| Nifedipine | C<sub>17</sub>H<sub>18</sub>N<sub>2</sub>O<sub>6</sub> | 389.1089 | 8.169 | 1.71E-09 | 8.20025<br>3 | 0.020819 |
| N-[2-(5-Ethyl-4-hydroxy-4,5-dimethyl-2-oxo-1,3-oxazolidin-3-yl)ethyl]-1-methyl-1H-pyrazole-5-carboxamide | C<sub>14</sub>H<sub>22</sub>N<sub>4</sub>O<sub>4</sub> | 371.1354 | 11.497 | 7.29E-10 | 7.95443<br>6 | 0.026787 |
| D-(+)-Galactose | C<sub>6</sub>H<sub>12</sub>O<sub>6</sub> | 535.1827 | 12.978 | 1.82E-09 | 7.692936 | 0.025431 |
| 10-Butyl-9,10-dihydroxy-4-pentyl-5,9,10,11-tetrahydro-4H-2,7-dioxadicyclopenta[a,e]cyclononene-1,3,6,8-tetraone | C<sub>22</sub>H<sub>28</sub>O<sub>8</sub> | 465.1772 | 17.607 | -7.38797 | 3.13E-06 | 3.72E-05 |

### Screening and Verification of Potential AR Therapeutic Components

#### Screening Metabolites with Potential Therapeutic Effects from SDO

When p value < 0.05, metabolites with log2FC > 1 or VIP > 1(log2FC > 0) and having credible names were integrated as the dataset to be analyzed. Due to the matching results of the mzCloud database being more reliable than the other two databases, the components related to AR treatment were screened among all metabolites with "full match" in the mzCloud database and other metabolites with log2FC > 3. After reviewed literature, a total of 11 compounds directly or indirectly related to AR treatment were found, they are DL-Arginine, ferulic acid, equol, nikethamide 1-oxide, 4-phenylbutyric acid, crotetamide, allantoin, nodakenin, 4-methyl-5-thiazolethanol, 4-coumaric acid, urocanic acid, and azelaic acid. Seven of them (DL-Arginine, ferulic acid, equol, nikethamide 1-oxide, 4-phenylbutyric acid, allantoin, nodakenin) are directly associated with AR treatment, while four (4-methyl-5-thiazolethanol, 4-coumaric acid, urocanic acid, and azelaic acid) are associated with anti-hypersensitivity or anti-inflammatory.^30,31,32,33,34,35,36,37,38,39^ Thus, 11 compounds were considered to have AR therapeutic potential.

Caused by the limitation of purchasement, 10 related compounds except for nikethamide-1were purchased for subsequent investigation. Furthermore, because SDO is a type of plant, the DL-Arg should be L-Arg in the original sample, so L-Arg was purchased for identification.

#### Metabolite Validations

First, QTOF-MS was performed on 10 standard references of metabolites with AR therapeutic potential to obtain the mass charge ratio (m/z), retention time (rt), and EIC images of each metabolite. Then, SDO and CMO water extract samples were subjected to QTOF-MS to obtain the EIC diagrams of samples. The corresponding chromatograms (EIC diagrams) were extracted based on the mz of each standard in the TIC diagrams of SDO and CMO water extract, and compared the peak with the corresponding standard on rt. When the rt of the extracted peak in the sample is consistent with that of the standard sample, it indicates that the compound has been verified to exist.

According to Figures 6 and 7, the DL-Arginine, urocanic acid, 4-methyl-5-thiazoleethanol, allantoin and 4-phenylbutyric acid were identified in the positive ionization results of QTOF-MS, while allantoin and azelaic acid were identified in the negative ionization results. However, the 4-methyl-5-thazoleethanol was only successfully identified in SDO water extract samples.

**Figure 6.**
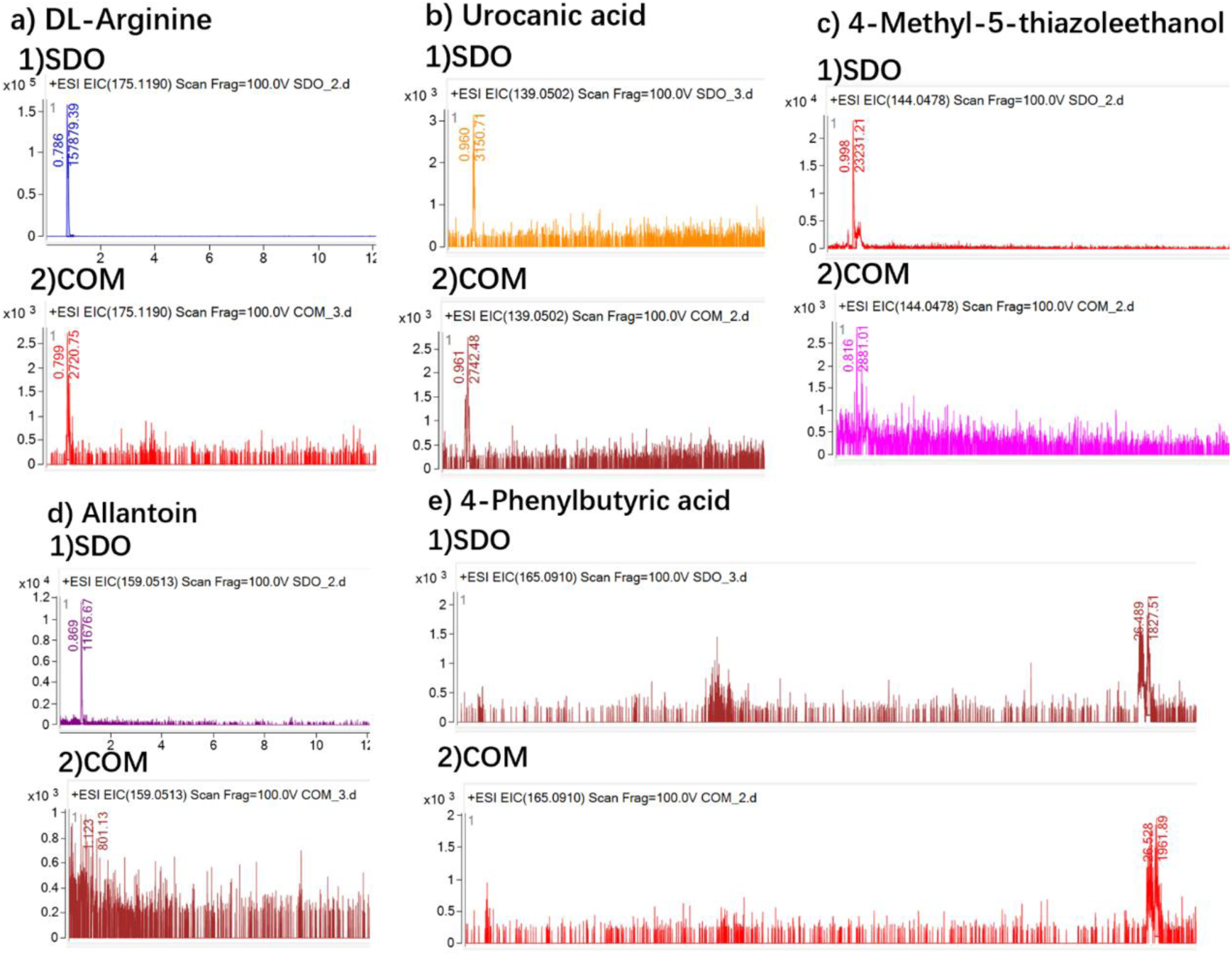
EIC diagrams of metabolites successfully verified in SDO sample at the positive ionization forms. SDO means the EIC diagrams is extract from the TIC result of the SDO sample, and COM means being extracted from COM sample’s TIC diagram. The number to the left of the peak is rt, and the number to the right is the peak area.

**Figure 7.**
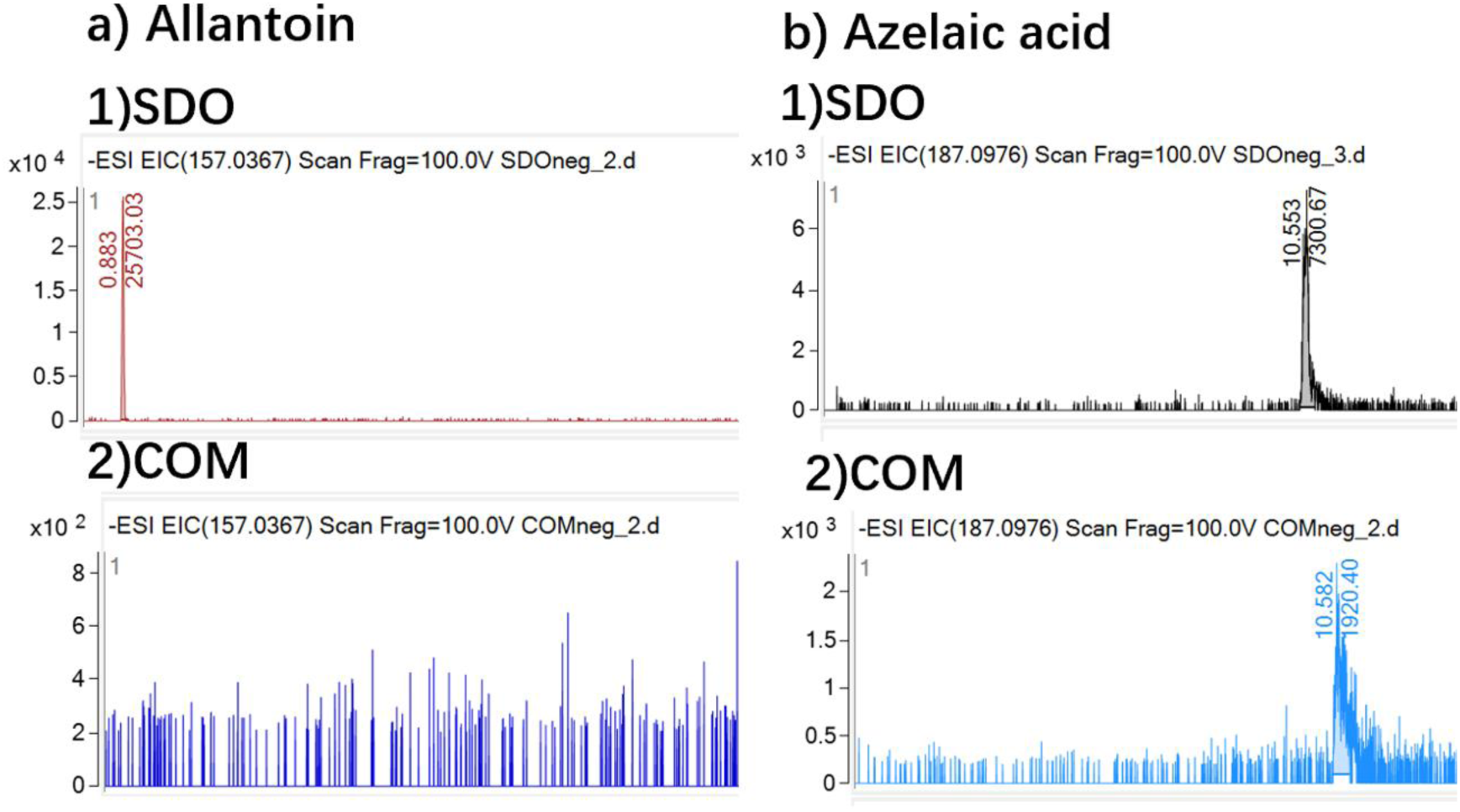
EIC diagrams of metabolites successfully verified in SDO sample at the negative ionization forms. The interpretation method is consistent with Figure 6.

Therefore, in these 10 compounds with potential AR therapeutic effects, a total of 6 compounds had been successfully identified, and allantoin appeared in the result of both ionization forms. However, compared with the rt of the HPLC-MS/MS identification result, the rt of 4-Phenylbutyric acid in the QTOF-MS result is different, so the 4-Phenylbutyric acid in the HPLC-MS/MS dataset may have been incorrectly identified.

### Network Pharmacological Analysis

#### Therapeutic Targets Prediction

Firstly, the information of six compounds with AR treatment potential obtained through the PubChem database was input into three drug target databases (PharmaMapper, Similarity Ensemble Approach, and Swiss Target Prediction database) to match their potential targets in the human body. To increase the relative credibility of the matching results, the top 100 targets from the PharmaMapper database, targets with p (probability) greater than 0 in the matching results of the Swiss Target Prediction database, and targets whose MaxTc is no less than 0.4 in the Similarity Ensemble Approach matching results were screened and merged into the drug target dataset. After removing duplicate matching results, a total of 255 potential targets were integrated into the drug target dataset.

Then, searching "Allergic rhinitis" in OMIM, Pharm GKB, TTD, Drug Bank, and Gene Cards databases found 2282 individuals’ AR targets and integrated into the AR target dataset (158 duplicate results were removed).

R language was used to screen potential AR therapeutic targets for the SDO water extracts that appear simultaneously in two datasets. According to the Venn diagram (Figure 8), 121 potential therapeutic targets were found.

**Figure 8.**
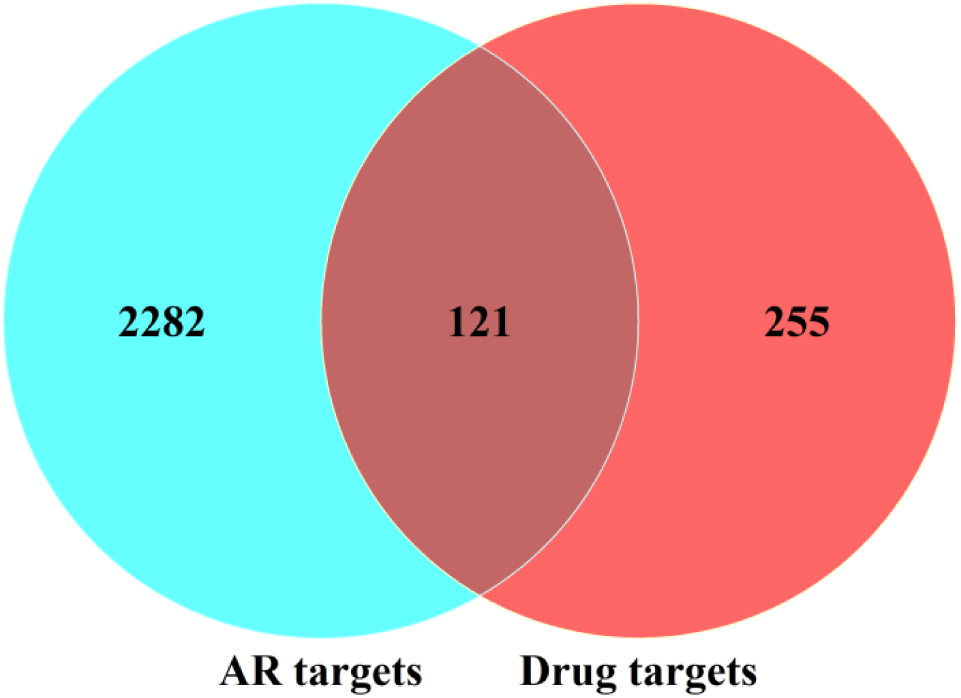
the Venn diagram of AR targets and drug (the 6 metabolites related to AR treatment) targets. The numbers on the figure are the number of targets.

#### Construction of the PPI network based on potential therapeutic targets

The 121 potential targets that may be related to SDO water extract treats AR were uploaded to the STRING database to search for the protein-protein interactions between targets and obtain the protein-protein interactions (PPI) network. Due to the number of potential targets, in order to obtain more valuable and reliable PPIs, the interaction score that was the minimum required was set to 0.7 (high confidence). Figure 9 shows the image of the generated PPI network, consisting of 279 edges and 121 nodes. The p-value of the protein-protein interactions enrichment from the PPI network is less than 1.0e-16, which indicates the existence of biological connections between the potential target proteins. The short tabular text of the PPI network is output as a TSV file for further topological analysis to determine the core target.

**Figure 9.**
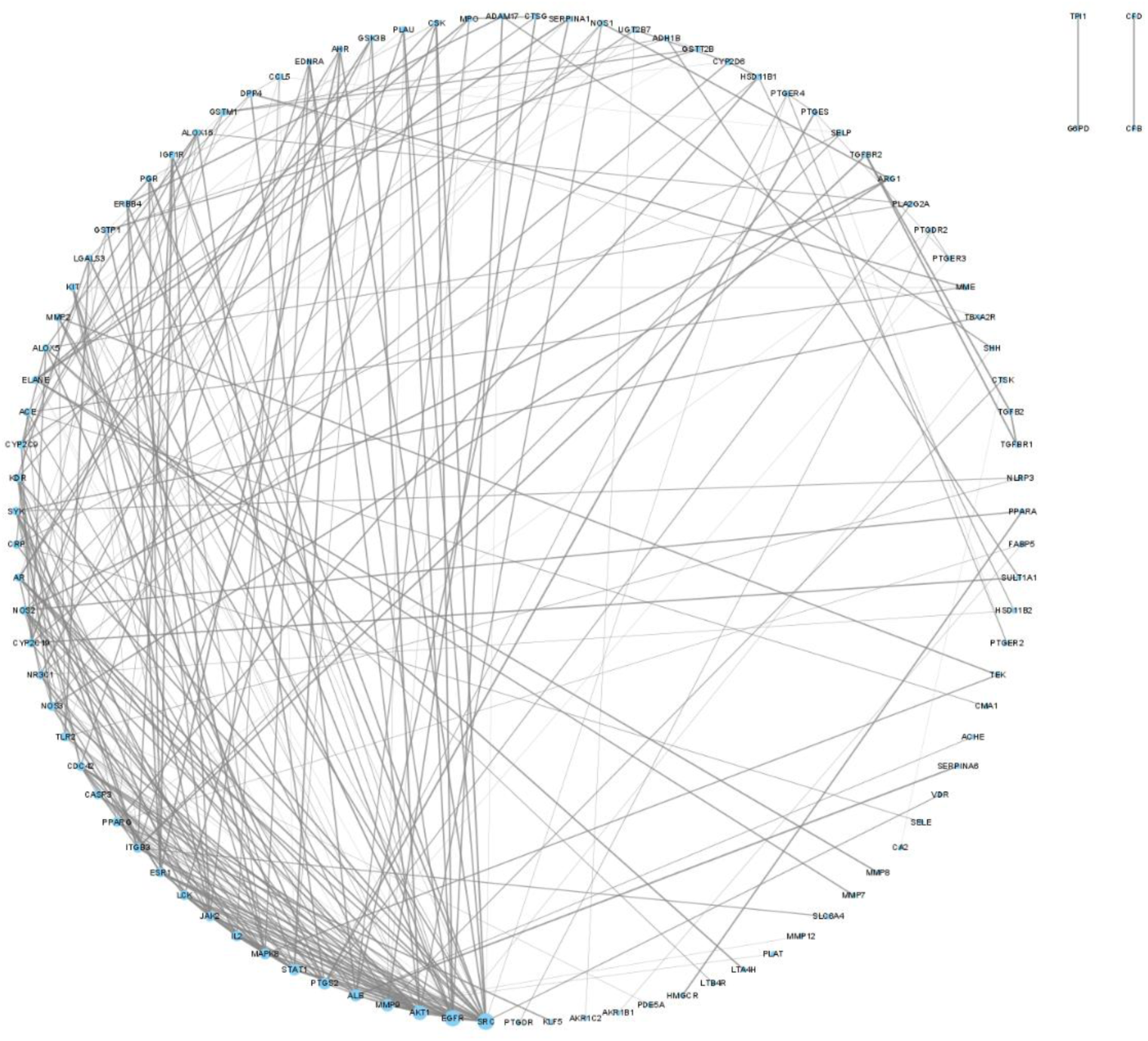
the PPI networks of the potential therapeutic targets Each point represents a target, and the line represents the interaction between the targets.

#### Screening of the potential core therapeutic targets

The TSV files of the PPI network constructed from 121 potential therapeutic targets were input into Cytoscape 3.9.1 to screen the core targets according to the network topological structure using the cytoHubba plug-in. According to the instruction of the cytoHubba plug-in, the algorithm based on maximum climate centrality (MCC) is the most effective in finding core nodes in this plug-in.^27^ Therefore, the MCC algorithm was used to calculate the scores of each node and screen the top 10 nodes with the highest scores as the potential core therapeutic targets. According to the descending order of scores, the top ten targets are EGFR, AKT1, SRC, MMP9, STAT1, ALB, LCK, CASP3, PTGS2, and IL2. Figure 10 shows the topological structure of these ten core targets in the PPI network.

**Figure 10.**
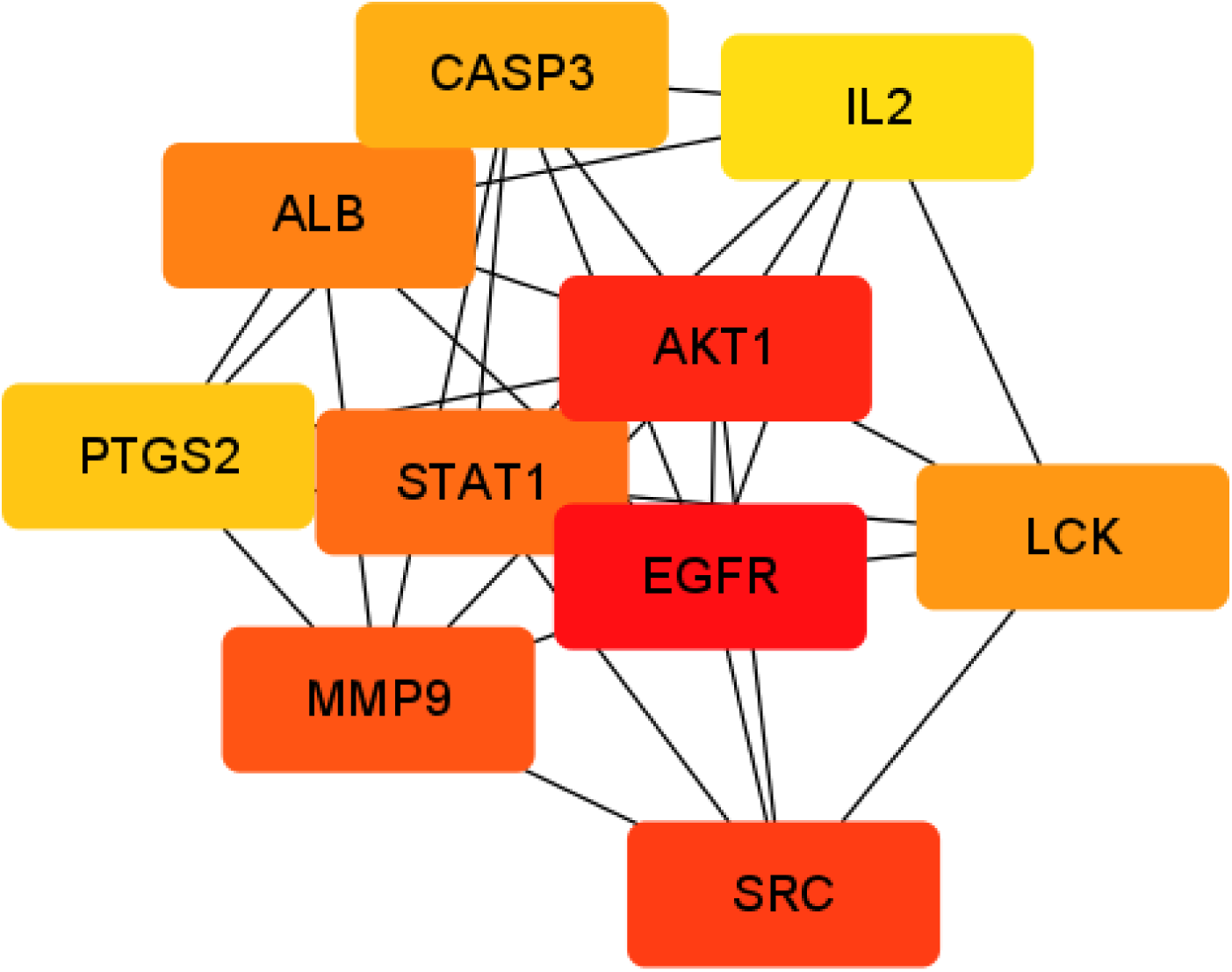
Top ten core targets obtained through the topology analysis The darker the target colour, the higher the MCC score of the target.

#### Enrichment analysis based on potential therapeutic targets

The 121 targets obtained by intersection of the drug and AR targets datasets were imported into R studio and converted into ENTREZID using the function in the R package "clusterProfiler".^40^ Subsequently, the obtained ENTREZID set was used for KEGG pathway and GO enrichment analysis by the R package "org. Hs. eg. db" and "clusterProfiler". The Benjamin Hochberg method (BH) was performed to correct the p-value of each GO term or pathway. When the p.adjust is less than 0.01, according to the descending order of gene ratio, the top 10 terms in each GO category (molecular function, MF; biological process, BP; cell component, CC) and the top 20 terms in KEGG pathway analysis results were selected as the main (Figure 11).

**Figure 11.**
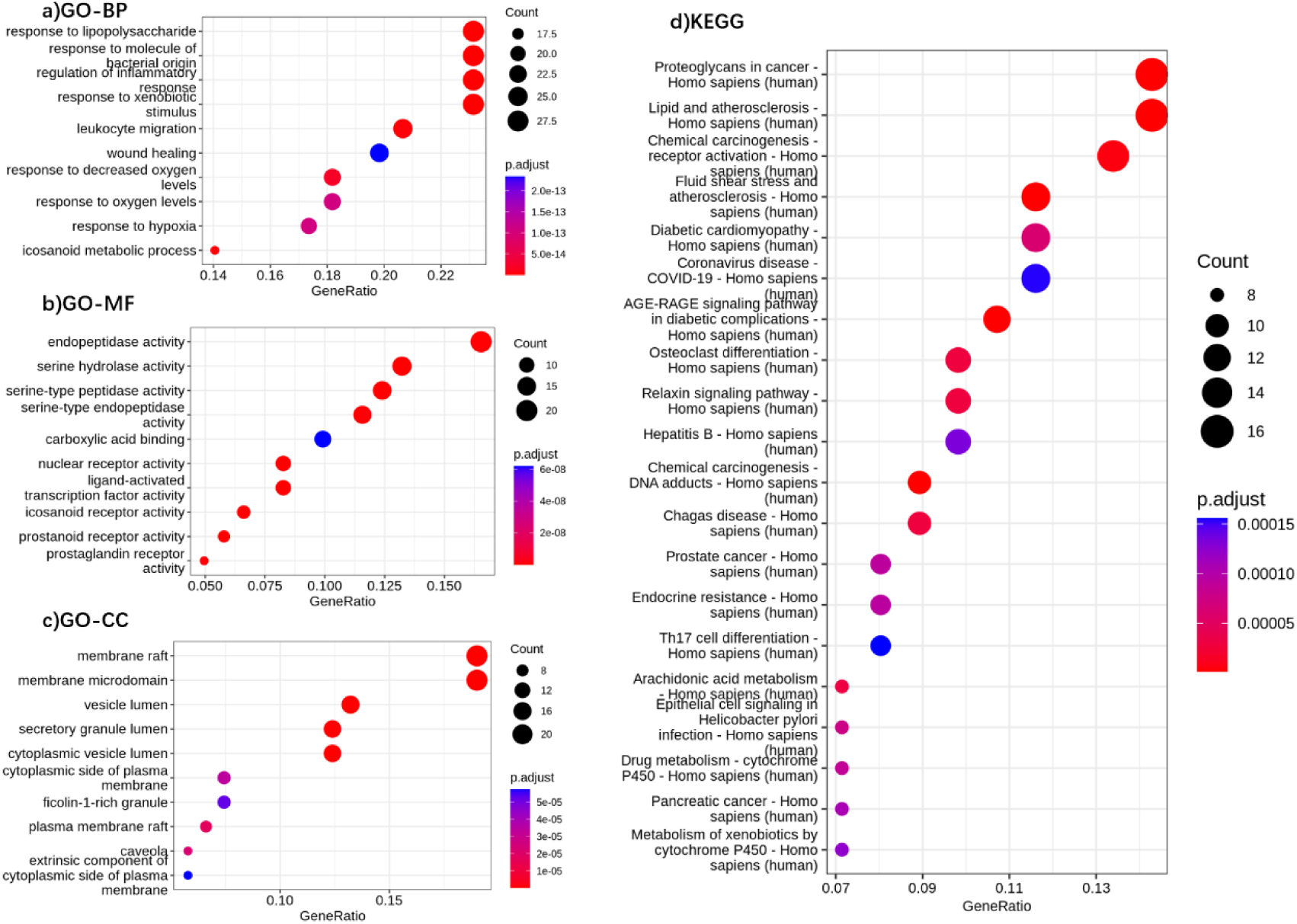
Enrichment analysis results. a,b,c: Top 10 GO enrichment terms in each category. d: Top 20 KEGG pathways based on the gene targets. p. Adjust is the p-value corrected by BH. Count is the number of genes in this term/pathway in the target list. Gene ratio is the gene numbers in this term/pathway: the number of genes annotated by KEGG/GO in the target gene set.

According to the figure 11, the top four terms in the BP class term have the same gene ratio values, they are "regulation of inflammatory response", "response to lipopolysaccharide", "response to xenobiotic stimulus", and "response to molecule of bacterial stimulus". Among the top 10 BP terms, the terms directly related to AR include "regulation of inflammatory response", "response to lipopolysaccharide", "leucocyte migration", and "icosanoid metabolic process".^41,42,43^ In the enrichment analysis results of MF terms, the three terms with the highest gene ratio are "endopeptidase activity", "serine hydroxylase activity", and "serine type peptidase activity". The "icosanoid receptor activity" and "prostanoid/ prostaglandin receptor activity" are directly related to AR in the top ten terms of MF.^43^ In CC terms, "member raft", "member microdomain", and "vessel lumen" are the three main terms involved (the top three terms according to the gene ratio), while the top ten CC terms with the highest enrichment are not directly related to AR.

According to the results of KEGG pathway analysis, the three metabolic pathways with the highest enrichment are "lipid and atherosclerosis", "fluid shear stress and atherosclerosis" and "osteoclast differentiation". Among the top 20 pathways with the highest gene ratio obtained from KEGG pathway analysis, "Th17 cell differentiation" and "arachidonic acid metabolism" are metabolic pathways related to AR.^44,45^

#### Molecular docking

The most expected binding energy between the top ten core target and 6 metabolites related to AR treatment is evaluated by molecular docking (Figure 12). The lower the binding energy, the higher the docking affinity, and the more stable the binding between the target and the compound. Each target has a binding energy of less than -5 kcal/mol with at least one compound, which indicated good binding ability. Among all the docking configurations, ALB has the lowest binding energy (-7.50 kcal/mol) with 4-phenylbutyric acid. The binding energies of 4-phenylbutyric acid and all core targets is less than -5 kcal/mol, indicating that it has good docking potential with these ten core targets. However, no core targets has binding energy less than -5 kcal/mol with 4-Methyl-5-thiazoleethanol, which may mean that the optimal AR therapeutic target of 4-Methyl-5-thiazoleethanol is not among these ten core targets.

**Figure 12.**
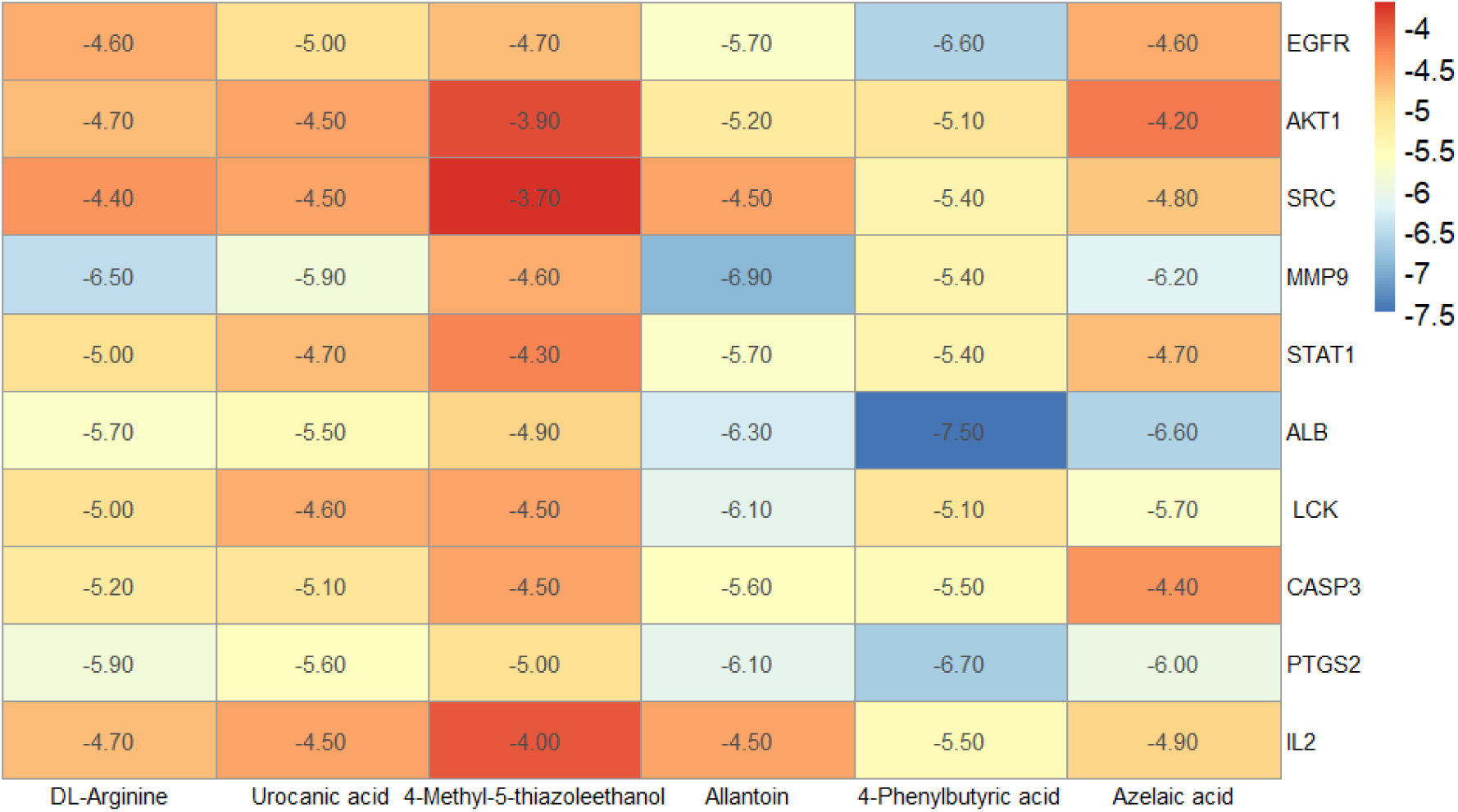
Heatmap of binding energy between top 10 core targets and 6 metabolites related to AR treatment. The numbers on the figure represent the minimum binding energy (Kcal/mol) in docking.

For concreteness, the docking configurations with binding energies less than -6.5 kcal/mol are shown in Figure 13.

**Figure 13.**
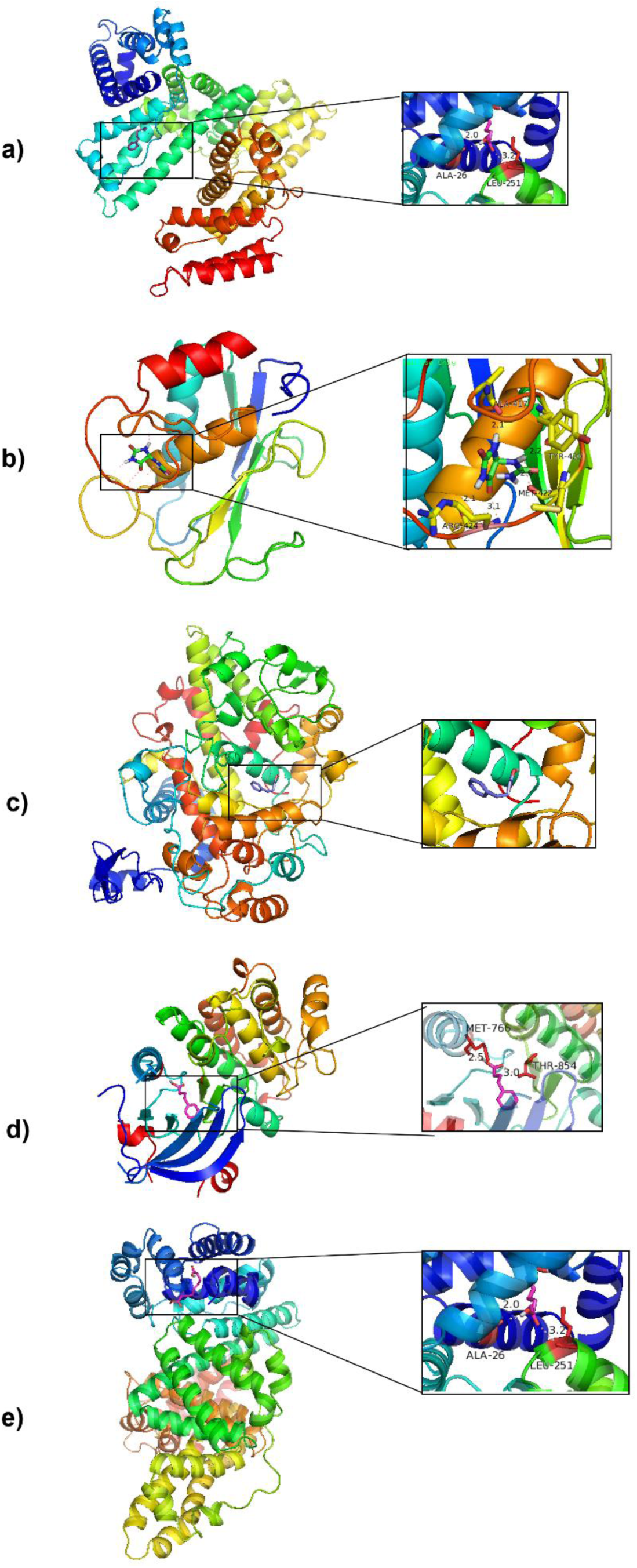
Molecular docking models with binding energies < -6.5 kcal/mol. a: Docking sites of ALB to 4-phenylbutyric acid. b: MMP9 to allantoin. c: PTGS2 to 4-phenylbutyric acid. d: EGFR to 4-phenylbutyric acid. e: ALB to azelaic acid.

## Discussion

As one of the post genomics’ cornerstones, metabolomics could conduct quantitative analysis and identification of small molecules fairly, and provide non-invasive global analysis for the biomarkers.^46^ HPLC-MS/MS based metabolomics can be performed to comprehensively understand herbal medicines’ chemical diversity and pharmacological characteristics.^1,46^ This study used HPLC-MS/MS based metabolomics to perform the metabolomics differential analysis on the metabolomics of SDO and COM water extracts. OPLS-DA analysis shows significant differences in the metabolomics of SDO and COM water extracts at the macro level, so the metabolome of SDO cannot be simply simulated using COM metabolome that already have partial results analysis and identification. Subsequent differential analysis of specific metabolites identified 2053 compounds with differential expression between the two water extracts. In these compounds,175 metabolites were relatively upregulated in SDO water extracts relative to the COM water extract, and 1878 metabolites were downregulated. Therefore, a small number of metabolites may be more enriched in expressing of the SDO metabolome.

According to the result of differential analysis, 10 metabolites related to AR treatment were screened and validated among the more reliable metabolites with higher fold change and expression differences between the two groups. Firstly, DL-Arginine, urocanic acid, 4-methyl-5-thazoleethanol, allantoin, 4-phenylbutyric acid and azelaic acid (6 metabolites) were successfully identified in the SDO water extract sample. These metabolites may belong to the core metabolites of SDO in treating AR. But even if metabolites with relatively reliable identification results were selected for validation, only six of the ten metabolites were successfully validated, and even one of them was identified incorrectly by HPLC-MS/MS. A fact that even performing strict quality control on results from HPLC-MS/MS, the obtained metabolomic results are not entirely reliable and need to be validated. Due to the time and funding limit, this study only screened a small number of metabolites with expression differences (and SDO > COM) and related to AR treatment for identification. Therefore, the bioactive substances that had relatively higher expression levels but no expression differences were ignored in screening, but they may also relate to AR treatment. In addition, according to previous studies, polysaccharides are the main bioactive components of Dendrobium Officinale stem, but due to their solubility, it is difficult to identify the polysaccharides in water extracts.^47^ Thus, it may be important to further comprehensively identify the bioactive components in SDO in further study, to specifically study the therapeutic mechanism of SDO on AR or other diseases. Secondly, only DL-Arginine, urocanic acid, allantoin, 4-phenylbutyric acid and azelaic acid were successfully validated in COM water extract samples. Nevertheless, the 4-methyl-5-thazoleethanol validation failed, which different from the SDO water extract. It may indicate that there have differences in efficacy and mechanism between SDO and COM water extract in treating AR.

Network pharmacology can explore the potential mechanisms in the synergistic therapeutic phenomenon of multiple drug molecules, and is the forefront of drug research and development.^48^ Due to network pharmacology’s characteristics, it is often used to study the therapeutic mechanisms of herbs for diseases.^49^ This study screened potential therapeutic targets and AR targets for AR therapy based on the 6 successfully identified AR therapy related metabolites from SDO water extract. Then, it further obtained 121 potential therapeutic targets that SDO water extracts treat AR by finding intersections. The protein-protein interaction between these 121 targets were plotted as a PPI network. The researcher adapted MCC algorithm, and found the 10 most important targets based on the topology of the PPI network. These targets are EGFR, AKT1, SRC, MMP9, STAT1, ALB, LCK, CASP3, PTGS2 and IL2. Molecular docking proved that the core targets combined well with the validated bioactive ingredients. EGFR can influence the secretion of PDGF, IL-8, and mucins by nasal epithelial cells.^50^ AKT1 plays a crucial role in cell apoptosis and inflammation, and may enhance the activation of NF-κB.^50^ MMP9 can regulate the release of cytokines and inflammatory factors, the increase of it was proved to further activate mast cells.^51^ STAT1 has a crucial role in Th1 lymphocyte differentiation, and it mediates the airway structural and immune cells’ pro-inflammatory response.^52^ LCK is highly correlated with signal transduction in inflammation and immunity.^53^ CASP3 may participate in cellular immunity through the TNF signaling pathway.^54^ PTGS2 mediates the eosinophil leukocyte migration in the inflammatory response.^55^ IL-2 is a pleiotropic cytokine that plays a crucial role in T cell metabolic and transcriptional programs.^56^

The results of KEGG pathway analysis and GO enrichment analysis based on 121 potential therapeutic targets show that the therapeutic effect of SDO aqueous extract on AR may be mediated by the following mechanisms: inflammatory response regulation, leucocyte migration, response to lipopolysaccharide, the metabolic process and receptor activity of icosanoid (i.e.

Eicosanoids, which include prostanoid/ prostaglandin), the differentiation of Th17 cells and arachidonic acid metabolism. Among the ten core targets, seven of them are directly related to inflammation regulation, so the therapeutic mechanism of SDO water extract on AR most probably be the multi-target mediated inflammatory response regulation.

The study has two limitations. First, it only used a small number of identified metabolites for target screening, the potential therapeutic targets of metabolites that are effective in AR treatment but have not been identified may be ignored. Therefore, the obtained network pharmacology analysis results may not be comprehensive enough. In addition, because network pharmacology results were only generated by computer simulations, the results had value for reference but were not entirely reliable. In subsequent studies, to confirm the specific molecular mechanism of SDO water extract in treating AR, more metabolites with high content will be identified and analyzed, and wet-lab experiments will be performed to validate the selected core targets and potential therapeutic mechanisms.

## Conclusion

In conclusion, this study combines metabolomics and network pharmacology to explore the medical potential of SDO for the first time. After verifying the therapeutic effect of SDO on AR through the preliminary experiment, the researcher performed metabolomics differential analysis on the water extracts of SDO and COM, identified some differential metabolites related to AR treatment, and inferred the core targets and therapeutic mechanisms involved in SDO treating AR. It provided a promising direction for the large-scale medical application of Dendrobium Officinale sprouts. However, there is still space for further improvement and research in the quantity and method of differential metabolites identification, and the exploration and validation of therapeutic mechanisms.

## Abbreviations

AR: Allergic rhinitis
SDO: the stem cell-originated Dendrobium Officinale
COM: commercial Dendrobium officinale stem
SVR: support vector regression
PPI: protein-protein interaction
MF: molecular function
BP: biological process
CC: cell component

## Acknowledgments

Sincerely thanks to my other supervisor Professor Cheng Ken for her invaluable guidance throughout my research. Furthermore, thanks to Suzhou Henghui Testing Co., Ltd. and Suzhou Institute of Nanobiology, Chinese Academy of Sciences for testing technology support.

